# Effects of ozone on sickness and depressive-like behavioral and biochemical phenotypes and their relations to serum amyloid A and kynurenine in blood

**DOI:** 10.1101/2021.12.16.472859

**Authors:** Kristen K. Baumann, W. Sandy Liang, Daniel V. Quaranta, Miranda L. Wilson, Helina S. Asrat, Jarl A. Thysell, Angelo V. Sarchi, William A. Banks, Michelle A. Erickson

**Author notes:** Corresponding Author: Dr. Michelle A. Erickson, Mailing Address: 1660 S. Columbian Way S-182, Seattle, WA, 98108.

## Abstract

Ozone (O_3_) is an air pollutant which primarily damages the lungs, but growing evidence supports that O_3_ exposure can also affect the brain. Serum amyloid A (SAA) and kynurenine have been identified as circulating factors that are upregulated by O_3_, and both can contribute to depressive-like behaviors in mice. However, little is known about the relations of O_3_ exposure to sickness and depressive-like behaviors in experimental settings. In this study, we evaluated O_3_ dose-, time- and sex-dependent changes in circulating SAA in context of pulmonary inflammation and damage, sickness and depressive-like behavioral changes, and systemic changes in kynurenine and indoleamine 2,3-dioxygenase (IDO), an enzyme that regulates kynurenine production and contributes to inflammation-induced depressive-like behaviors. Our results in Balb/c and CD-1 mice showed that 3ppm O_3_, but not 2 or 1ppm O_3_, caused elevations in serum SAA and pulmonary neutrophils, and these responses resolved by 48 hours. Sickness and depressive-like behaviors were observed at all O_3_ doses (1-3ppm), although the detection of certain behavioral changes varied by dose. We also found that *Ido1* mRNA expression was increased in the brain and spleen 24 hours after 3ppm O_3_, and that kynurenine was increased in blood. Together, these findings indicate that acute O_3_ exposure induces transient symptoms of sickness and depressive-like behaviors which may occur in the presence or absence of overt pulmonary neutrophilia and systemic increases of SAA. We also present evidence that the IDO/kynurenine pathway is upregulated systemically following an acute exposure to O_3_ in mice.

## Introduction

Ozone (O_3_) is a widespread toxicant in air pollution that is harmful to human health. Epidemiological studies have shown that short-term increases in ambient O_3_ are associated with increased mortality and morbidity ^1-3^. Healthy humans acutely exposed to O_3_ experience temporary adverse effects, which include decreased pulmonary function and increased pulmonary inflammation ^4-6^. Although the lungs are the primary target organ for direct oxidant activities of O_3_, growing evidence supports that O_3_ exposure may affect distal organs, including the brain ^7,8^. Increased oxidative stress, neuroinflammation, blood-brain barrier (BBB) disruption, and neurotransmitter dysfunction have been associated with O_3_ exposure in rodents and humans ^9-13^. Epidemiological studies also implicate O_3_ as a possible risk factor for neurological conditions such as cognitive impairment/decline ^14-16^, Alzheimer’s disease (AD) ^17^, and depression ^18-21^ in certain populations. Acute exposures to O_3_ can elicit neurological symptoms such as headache, dizziness, fatigue, and mental tension in humans ^22,23^, suggesting that CNS effects of O_3_ may contribute to exposure-associated morbidities such as feelings of illness in the short-term. However, the mechanisms by which O_3_ elicits its effects on the CNS are largely unclear, since the direct oxidant activities of O_3_ are limited to the lungs

One emerging hypothesis on how O_3_ affects the CNS is through a lung-brain axis mechanism ^12^ whereby initial O_3_-induced pulmonary damage and local inflammation elicit a systemic inflammatory response and the release of circulating factors that can affect CNS functions. Often, neuroimmune communication pathways involve the systemic release of pro-inflammatory cytokines and chemokines ^24^, but blood and brain levels of pro-inflammatory cytokines and chemokines are not increased in mice exposed to O_3_ ^12,25^. However, other inflammation-associated factors such as kynurenine and serum amyloid A (SAA) are increased in the blood of O_3_ -exposed in rats and mice, respectively ^25,26^. Kynurenine is a tryptophan metabolite that is elevated in blood and brain in response to inflammation, through the activation of its rate-limiting enzyme, indoleamine 2,3-dioxygenase (IDO) ^27^. Activation of IDO has been shown to mediate depressive-like behaviors following inflammatory stimuli ^28,29^, and kynurenine elevations have been implicated in depression ^30^ and in other neurological diseases ^31^. Kynurenine can cross the intact BBB via the large neutral amino acid transporter, LAT-1, and it has been estimated that most of the kynurenine in brain is derived from the circulation ^32^. SAA is an acute phase protein and two of its isoforms, SAA1 and SAA2, are produced mainly by hepatocytes, are markedly upregulated in response to inflammatory insults, and circulate as HDL lipoproteins ^33^. We recently showed that SAA1 and SAA2 can cross the intact mouse BBB, and that SAA1/2 concentrations in brain, liver, and blood increase following an acute O_3_ exposure in mice ^25^. In the same study, SAA levels in blood correlated significantly with O_3_-induced pulmonary inflammation. Elevations of SAA in blood have also been associated with pulmonary inflammation during acute exacerbations of COPD ^34^, suggesting that pulmonary inflammation is coupled with SAA production by the liver in human disease.

The current evidence that both SAA and kynurenine are induced by O_3_, derive in part from peripheral sources and can cross the BBB, contribute to depressive-like behaviors in mice ^29,35^, and are associated with depressive symptoms in humans ^30,36^ suggests that they are important mediators of neurobehavioral changes following O_3_ exposure. Therefore, we sought to further characterize the O_3_- induced SAA and kynurenine increases and their relation to pulmonary inflammation and acute changes in behaviors. Behaviors associated with sickness and depression can be overlapping and include malaise, anorexia, reduced locomotor activity, disinterest in social interactions, lethargy, reduced grooming, weight loss, anxiety, anhedonia, and memory impairment ^37,38^. Sickness behaviors are distinguished from depression, in part, by their manifestation as an adaptive response to infection or injury ^38^, however inflammatory challenges that initially cause sickness behaviors can also cause depressive symptoms. For example, pharmacological treatment with interferon-α (IFN-α) can precipitate a major depressive episode in 15-40% of patients ^39^. In humans experimentally treated with a low dose of bacterial lipopolysaccharide (LPS), symptoms of anxiety, depressed mood, and memory impairment are transiently induced and are more severe in women ^40,41^. In mice, a single low-dose injection of LPS can contribute to both sickness and depressive-like behaviors, which are expressed in distinct temporal patterns ^28,30^. Behaviors such as reduced sucrose preference and increased immobility in the tail suspension test persisted after other behaviors associated with cytokine-induced sickness responses including reduced food intake and locomotor activity resolved ^28^. To our knowledge, the temporal patterns of sickness and depressive-like behaviors as they relate to inflammatory changes following acute O_3_ exposure have not been evaluated in mice. However, such information would be important for the design of studies that evaluate depressive-like behaviors, cognition, or other behavioral phenotypes that could be influenced by acute behavioral responses related to sickness.

The first goal of our study was to evaluate the dose-and time-responses of sickness and depressive-like behavioral changes in female Balb/c and CD-1 mice, with respect to pulmonary inflammation and damage, and SAA levels in blood. We also aimed to determine whether sex influences any of our measured parameters. Our results show that there is a defined O_3_ dose-threshold and time-window that induces elevations in SAA, pulmonary inflammation, and sickness and depressive-like behaviors, and there are sex differences in these responses to O_3_. We further showed that tissue IDO expression and kynurenine levels in blood were significantly altered by O_3_, and that O_3_ induced similar biochemical and behavioral responses in female Balb/c and CD-1 mice, highlighting the robustness of the response across mouse strains.

## Materials and Methods

### Vertebrate animals

All mice were treated in accordance with NIH Guidelines for the Care and Use of Laboratory Animals in an AAALAC-accredited facility and approved by the Institutional Animal Care and Use Committee of the VA Puget Sound Health Care System. Male and female BALB/c mice and female CD-1 mice were purchased from Charles River, allowed to adapt for 1–2 weeks following shipment and were studied at 10-12 weeks of age. Mice were kept on a 12/12-h light/dark cycle (6:00-18:00 lights on) with ad libitum food and water, except during exposures to O_3_ when food was withheld. Power analysis was used to determine animal numbers needed for each study following initial pilot studies to estimate variance. A total of 201 mice were used in this study, and the numbers allocated to each group/figure are shown in Table 1.

**Table 1.** Animal numbers used in each study.

|  | **Air** | **1ppm** | **2ppm** | **3ppm** | **Figures** |
| --- | --- | --- | --- | --- | --- |
| **Balb/c dose response (female)** | 20 | 15 | N/A | 10 | 1 |
| **Balb/c time response (female)** | 10 | N/A | N/A | 30 (24, 28, 72 hrs) | 2, 3, 4, 5 |
| **Balb/c 1ppm food, fluid, and sucrose preference (female)** | 10 | 10 | N/A | N/A | 4, 5 |
| **Balb/c male vs. female** | 10 male/10 female | N/A | N/A | 10 male/10 female | 4, 5, 8, 9 |
| **Balb/c olfaction (female)** | 5 | N/A | N/A | 4 | 4 |
| **CD-1 dose response** | 10 | 9 | 9 | 9 | 6, 7 |

### Ozone exposures

Just prior to exposures, mice were group housed (n=3-4/cage) in standard mouse cages with wire tops that lack bedding. Individual mice were identified using an animal-safe marker. Food was withheld for the duration of exposure to prevent consumption of ozonated food, and water was provided ad libitum. Up to 4 cages at a time were placed in a 30”x 20”x 20” polypropylene chamber where O_3_ (3ppm, 2ppm, or 1ppm, chamber 1) or compressed dry air (chamber 2) was pumped into the chamber at equivalent rates. Because we only had two chambers, only one O_3_ dose was co-exposed with air on a given day. However, O_3_ exposures at each dose were replicated at least once to capture day-to-day variability, and results for each dose showed consistent trends. Males and females were co-exposed in the same chamber for studies that compared sex. Each chamber is equipped with a small fan near the infusion site that ensures even dispersion of the infused gas throughout the chamber. The temperature was maintained at 21-24 °C and the humidity at 35-49% for both chambers. O_3_ levels in the chambers were generated and regulated using an Oxycycler AT42 system (BioSpherix, Parish NY, USA). Prior to each experiment, the system was calibrated using a model 106-L O_3_ detector (2B Technologies, Boulder, CO, USA), and O_3_ levels were recorded from an inlet valve in one of the mouse cages every 10 seconds for the duration of exposures. In all studies, O_3_ achieved its target concentration within 10 minutes, and levels were regulated within 10% of the target concentration (1ppm +/- 0.1ppm and 3ppm +/- 0.3ppm) thereafter. All exposures were conducted for 4 hours (10:00-14:00), and mice were then returned to their home cages. Figure 1 depicts the set-up of our O_3_ exposure paradigm with respect to timing of behavioral assays and tissue collection. Behavioral testing and tissue collection was done in a randomized order so that each group was evenly dispersed across each time window of testing to mitigate nuisance variables.

**Figure 1:**
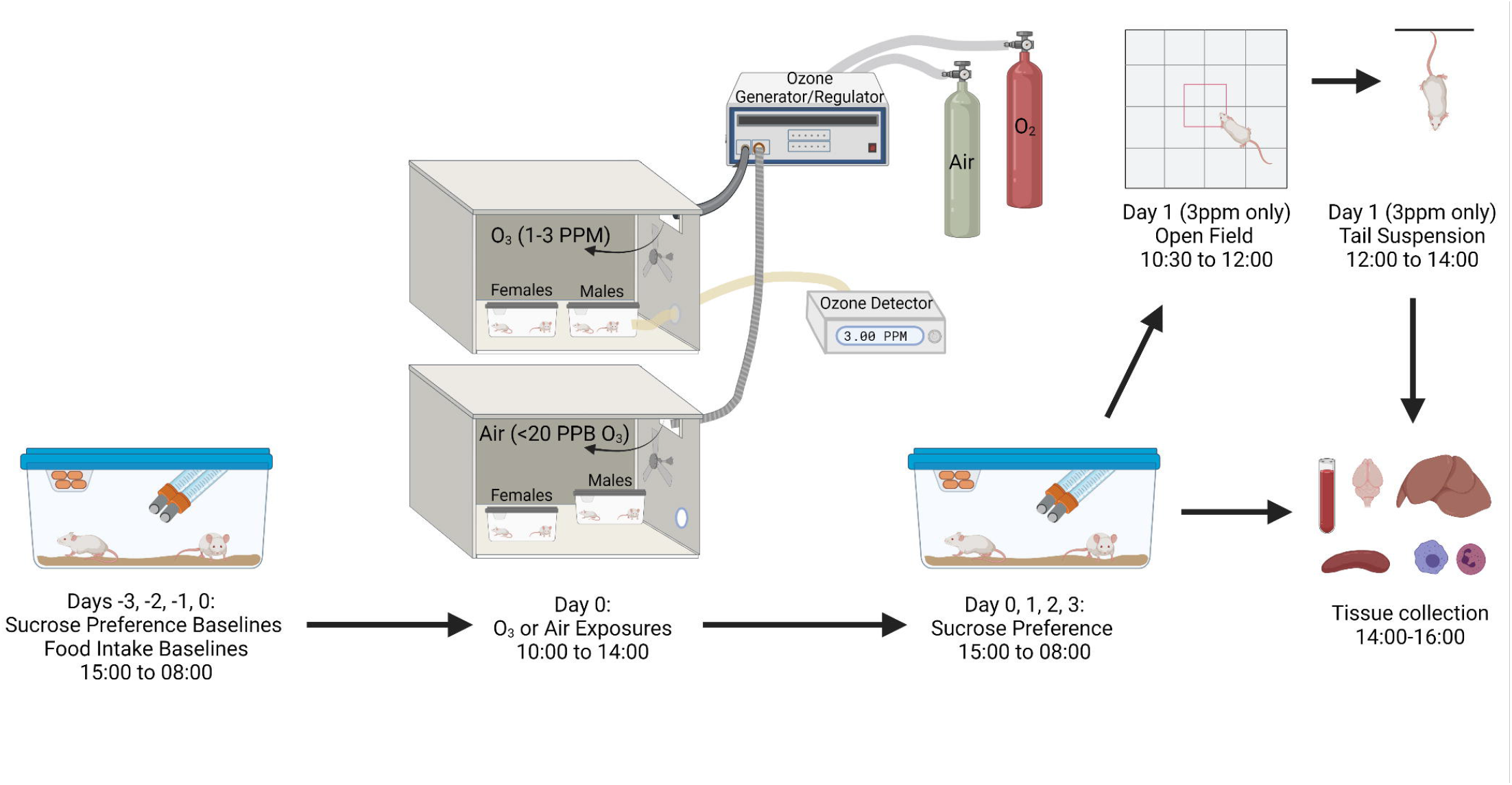
Schematic for the schedule of O_3_ exposures, behavior testing, and tissue collection

We have previously used an O_3_ exposure paradigm of 3ppm for 2 hours to characterize systemic changes in the acute phase protein SAA as they relate to airway inflammation ^25^. This paradigm optimally induces airway inflammation and obstruction without causing respiratory distress ^42^. When attempting to replicate these results using a different exposure apparatus at a different institution, and under group-housing conditions to mitigate stress that could affect results of behavioral studies, we found that a longer exposure time of 4 hours was needed to significantly induce increases in airway inflammation and SAA in serum to similar levels as observed previously ^25^. Although O_3_ concentrations that are environmentally relevant to humans are much lower than those used in this study, it has been shown that higher concentrations of O_3_ are needed to elicit similar pulmonary responses in experimental exposures of rodents vs. humans. For example, healthy young men exposed to an environmentally relevant dose of 0.1ppm O3 for 6.6 hours with moderate exercise exhibited over a 350% increase in BAL neutrophils, and over a 20% increase in BAL protein 18 hours post-exposure ^43^. Healthy young men exposed to 0.4ppm O_3_ for 2 hours with exercise had more robust responses, with neutrophils increasing over 8-fold and BAL protein increasing 2-fold ^44^. In our study, female BALB/c mice exposed to 1ppm O_3_ exhibited an 8.8-fold increase in BAL neutrophils (although this was not significantly different from controls), and a 0.5-fold increase in BAL protein. Therefore, the magnitude of biological responses observed at 1ppm in mice are similar to those reported in human studies using a short-duration 0.4ppm dose with intermittent exercise in healthy human males. This difference in humans and mice may be explained, in part, by less O_3_ deposition in the airway under typical rodent exposure conditions that do not involve exercise ^45^, as well as a greater resistance of rodents to O_3_ -induced damage and inflammation ^46^. The 3ppm dose elicits higher fold-changes in BAL and neutrophils, but we included this dose for comparison to our prior work for the purpose of comparing to lower doses of 2ppm and 1ppm O_3._

### Sucrose preference test

A decreased preference for sucrose as measured by this test is taken as evidence for depression, and sucrose preference testing was carried out based on standard methodologies that do not withhold food ^47^. Mice were single-housed and adapted to drinking water from two sipper bottles filled with water for 3 days. The morning after habituation, one of the water bottles was replaced with a 3% sucrose solution, which was determined to be the optimal concentration of sucrose to achieve about 80% preference on average for the Balb/c strain and 90% on average for the CD-1 strain. Overnight liquid consumption was then evaluated by weighing the bottles at 15:00 and again at 8:00. Baseline sucrose consumption was recorded for three nights, and each morning the bottles were switched in their left/right orientation. Sucrose preference testing post-ozone commenced at 15:00 the day of exposures and bottles were re-weighed at 8:00 the next day. Mice showing a baseline sucrose preference of 50% or lower (which we found occurs at about a 10% rate in Balb/c mice) were excluded from analysis. Mice were also occasionally excluded if there was evidence of leakage or drainage of either sipper bottle by the mouse.

### Open field test

In this test, decreased locomotor activity is taken as evidence for sickness behavior ^29^. At 8:30 the morning after O_3_ exposure, mice were placed in a quiet room with dim light (8-10 lux) and allowed to adapt for 2 hours. They were then placed into the center of a square open field with dimensions 40×40 cm and activity in the field was recorded for 10 minutes using ANY-maze video tracking software (Stoelting Co. Wood Dale, IL, USA). The center of the field was defined as a square with dimensions 20×20cm. The order of testing was randomized such that mice from each group were evenly distributed over the testing period.

### Tail suspension test

In this test, an increase in the time spent immobile vs. control is taken as evidence for depressive-like behavior. Experiments were carried out according to established methods for this test ^48^. Briefly, medical tape was placed around the mouse’s tail leaving some slack, and the slack of the tape was fastened using a binder clip. Thin, rigid plastic tubes were placed over the mouse tails to prevent tail climbing. The binder clip was hung on a hook, fastened to a 25cm (w) x 40cm (L) x 20cm (D) plastic box with the open end faced towards a camera. An empty mouse cage with a foam pad covered by absorbent bench paper was placed under the mouse in the event of a fall. Within about 10 seconds of securing the mouse, movement was tracked for 6 minutes using ANY-maze video tracking software (Stoelting Co. Wood Dale, IL, USA). The order of testing was randomized such that mice from each group were evenly distributed over the testing period.

### Olfaction test

As an impaired sense of smell can affect performance in the sucrose preference test, olfaction was tested in our mice using the buried food test ^49^. Mice were habituated to Froot Loop ® treats for three nights prior to study by providing about 5 per cage in the afternoon, and palatability of the treats was confirmed by verifying their consumption the next morning. Mice were then exposed to air or 3ppm O_3_ from 10:00-14:00, returned to home cages with food from 14:00-17:30, and fasted overnight until olfaction testing the next morning at 8:00. The next morning, mice were placed in a cage with about 3 inches of bedding that contained a buried Froot Loop ® treat in one of the corners. The mice were monitored and the time to find the treat was recorded. Due to the potential confounds of fasting, these mice were not included in assessments of other behavioral, physiological, or biochemical parameters.

### Tissue collection and processing

Mice were anesthetized with 2mg/g urethane, and blood was harvested from the abdominal aorta, allowed to clot at room temperature for 30 minutes, and then centrifuged at 2500xg for 10 minutes. Serum was collected, aliquoted and stored frozen. Brains were collected by severing the spinal cord, leaving the trachea intact for bronchoalveolar lavage, and then cut in half sagittally. The left medial lobe of the liver and spleen were also dissected and cut in half. One half of each tissue was snap-frozen in liquid nitrogen and the other half was placed in RNA later and frozen. Bronchoalveolar lavage (BAL) was performed on mice by perfusing and aspirating the lungs 3 times with 1 ml sterile phosphate buffered saline. BAL fluid was stored on ice and centrifuged at 200g for 5 minutes at 4 C. The supernatant was removed, aliquoted, and frozen and 0.5mls of supernatant was reserved to resuspend the cell pellet. Total cell counts were determined from the cell suspension by manual counting using a hemacytometer. Differential cell counts were performed on cytocentrifuge preparations (Cytospin 3, Thermo Fisher Scientific, Waltham, MA, USA) that were stained with Hemacolor (Sigma-Aldrich, St. Louis, MO, USA). Cell counts were performed using Image J, and at least 200 cells were counted to determine relative cell proportions. Exclusion of datapoints occasionally occurred if BAL was not recovered or if the cytospin slides were not countable.

### SAA ELISA

SAA mouse duoset kits (R and D systems, Minneapolis, MN, USA) were used to quantify SAA in serum. Serum was thawed and diluted 1/50 for controls and 1/10,000 for ozone-exposed samples using diluent specified in the kit. ELISAs were carried out according to manufacturer instructions. Some samples are missing because there was not sufficient serum recovered from some of the mice in the cohort.

### Kynurenine ELISA

IDK high sensitive Kynurenine ELISAs (Immundiagnostik, Bensheim, Germany) were used to quantify kynurenine in serum. Serums and standards were diluted and derivatized, and ELISAs carried out according to kit instructions. Some samples are missing because there was not sufficient serum recovered from some of the mice in the cohort.

### RNA extraction and IDO measurement

Tissues preserved in RNA later were thawed and homogenized in Qiazol. RNA was extracted from tissue homogenates using RNeasy Plus Universal Kits (Qiagen, Valencia, CA, USA). Superscript IV first strand kit (Thermo Fisher Scientific) was used to convert 5µg RNA template to cDNA in a 20ul reaction volume. For qPCR, 120 ng cDNA was amplified using TaqMan Fast Advanced Master Mix. TaqMan primer/probe sets with FAM-MGB probe (Thermo Fisher, Waltham, MA, USA) were used to amplify cDNA and included mouse *Ido1* (Mm00492590_m1), *Tdo2* (tdo2 Mm00451269_m1), and *Gapdh* (Mm99999915_g1). The delta-delta Ct method was used to calculate relative changes in *Ido1 and Tdo1* expression among treatment groups. Some samples are missing from spleen and liver mRNA measurements due to poor RNA quality following extraction.

### Statistical analysis

Analysis of dose- and time-response data with three or more groups was carried out by one-way ANOVA and individual groups compared using Tukey’s multiple comparisons test. Linear trends were also evaluated. Analysis of two groups was carried out by two-tailed unpaired T-tests. Analysis of sex-dependent responses to O_3_ was done using two-way ANOVA and individual groups were compared using Sidak’s multiple comparisons test. All data were analyzed using the Prism software package version 8.3.0 (GraphPad Inc, San Diego, CA).

## Results

### Dose effects of O_3_ on body weight, serum SAA, and pulmonary damage and inflammation

Our prior work demonstrated that female Balb/c mice exposed to 3ppm O_3_ had significantly increased levels of SAA protein in blood and brain 24 hours after exposure ^25^. We first wanted to determine whether SAA induction also occurred at lower concentrations of O_3_, and chose 1ppm as a lower dose because it has been used by others to evaluate CNS endpoints following acute exposure to O_3_ ^12,13^. We also evaluated weight loss and markers of pulmonary vascular leakage and inflammation in the same cohort of mice to understand how these measures relate to SAA changes in blood. All mice in this cohort were female. We found that mice exposed to 1ppm O_3_ did not show statistically significant increases in weight loss vs. air control, whereas mice exposed to 3ppm O_3_ in this cohort lost about 4.7% of their body weight, which was significantly different from the control and 1ppm groups (Figure 2A). 1ppm O_3_ caused no significant increase in serum SAA, with a mean concentration approximating that of air control. In contrast, 3ppm O_3_ increased serum SAA concentration by several orders of magnitude, consistent with what we have observed previously (Figure 2B) ^25^. Both 1ppm and 3ppm O_3_ significantly increased BAL total protein concentration, a marker of altered alveolar capillary barrier function ^50^, to similar levels that were not statistically different from each other (Figure 2C). 3ppm O_3_ significantly increased BAL total cells (Figure 2D) and BAL neutrophils (Figure 2F) vs. control, whereas 1ppm O_3_ did not. BAL macrophage counts were not significantly different among groups (Figure 2E). A summary of the effects of O_3_ dose in Balb/c mice is presented in Table 2.

**Figure 2:**
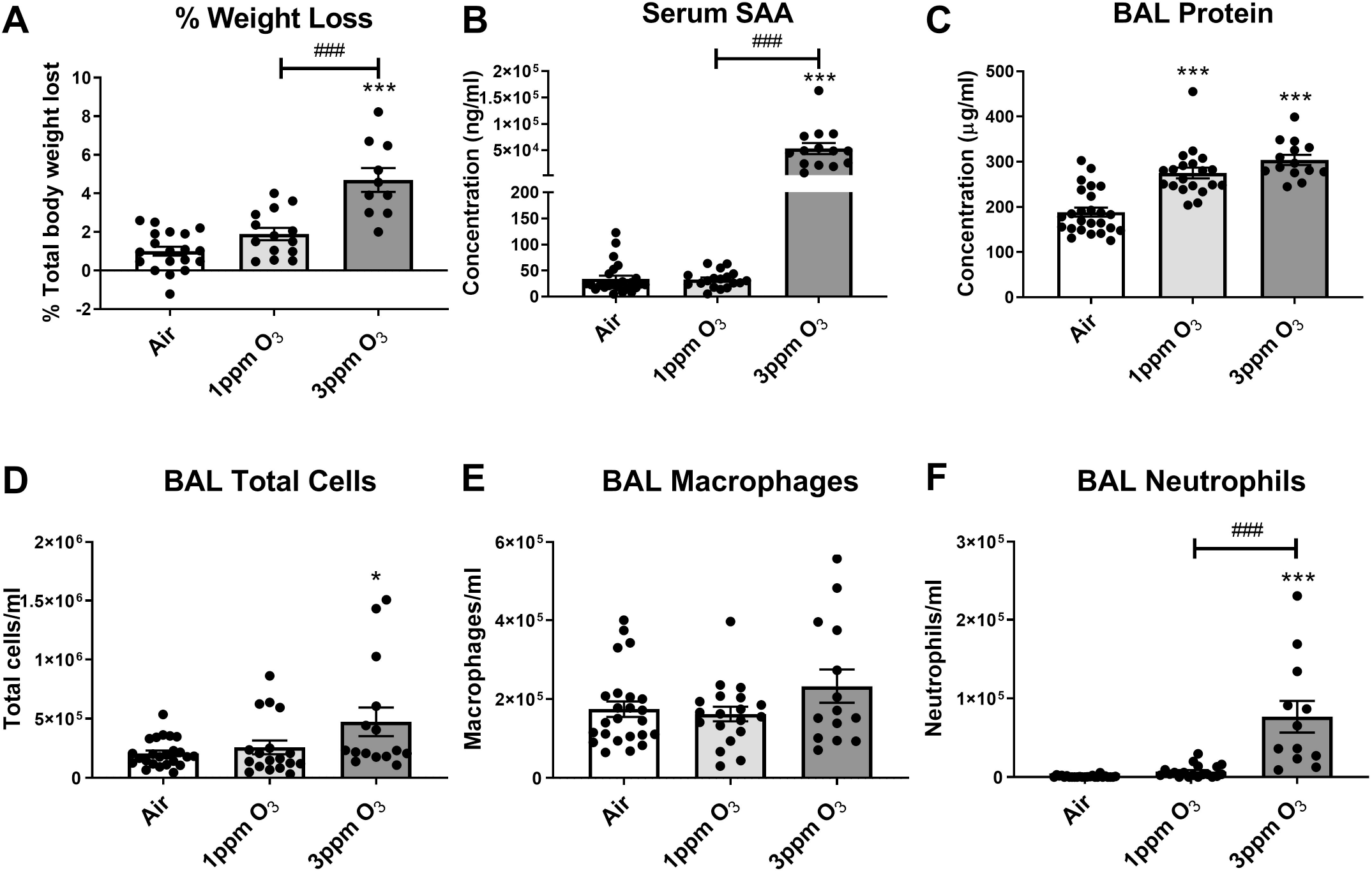
Effects of O_3_ concentration on weight loss (A), serum SAA (B), BAL total protein (C), and cellular markers of acute pulmonary inflammation (D-F). All mice were studied 24 hours post-exposure. N= 10-19/group, *p<0.05, ***p<0.001 vs. air, ###p<0.001 vs. indicated groups (1ppm vs. 3ppm).

**Table 2.** Summary of significant dose and sex-dependent changes induced by O_3_ in Balb/c mice. ↑ and ↓ indicate an increase or decrease vs. control, respectively. N.S. = no significant difference with O_3_, N.D. = not determined.

|  |  |  |  |  |
| --- | --- | --- | --- | --- |
|  |  | **1ppm O3** | **3ppm O3** | **Male vs. female** |

|  |  | (either sex) | (either sex) | difference |
| --- | --- | --- | --- | --- |
| **Pulmonary inflammation/damage markers** | Bal protein | ↑ | ↑ | No |
|  | Total BAL cells | N.S. | ↑ | Yes |
|  | BAL macrophages | N.S. | N.S. | Yes |
|  | BAL neutrophils | N.S. | ↑ | Yes |
| **Systemic inflammatory markers** | Serum SAA | N.S. | ↑ | Yes |
|  | Serum kynurenine | N.D. | ↑ | No |
|  | Brain *Ido1* | N.D. | ↑ | No |
|  | Spleen *Ido1* | N.D. | ↑ | No |
| **Sickness Behaviors** | Weight loss | N.S. | ↑ | Yes |
|  | Food intake | ↓ | ↓ | No |
|  | Fluid intake | ↓ | ↓ | No |
|  | Open field activity | N.D. | ↓ | No |
| **Depressive-like behaviors** | Sucrose preference | ↓ | ↓ | No |
|  | Tail suspension immobility | N.D. | N.S. | No |

Because of the apparent differences in responses to the 1ppm and 3ppm doses in female Balb/c mice, we next determined whether there was a similar dose effect relationship of measured parameters in female CD-1 mice. CD-1 mice were chosen because we wanted to replicate our experiments in an outbred strain as a more rigorous model. We also included an intermediate dose of 2ppm to have a more comprehensive understanding of the dose effects. Our findings in CD-1 mice supported a significant relation of O_3_ dose to the magnitude of changes in outcomes. Significant (p<0.001) linear trends were noted for weight lost (F (1, 33) = 25.62), total cells (F (1, 32) = 56.28), total neutrophils (F (1, 32) = 32.02), total macrophages (F (1, 32) = 48.74), BAL protein concentration (F (1, 31) = 24.51) and serum SAA (F (1, 33) = 25.62), and differences in group means are shown in Figure 3 and summarized in Table 3.

**Figure 3:**
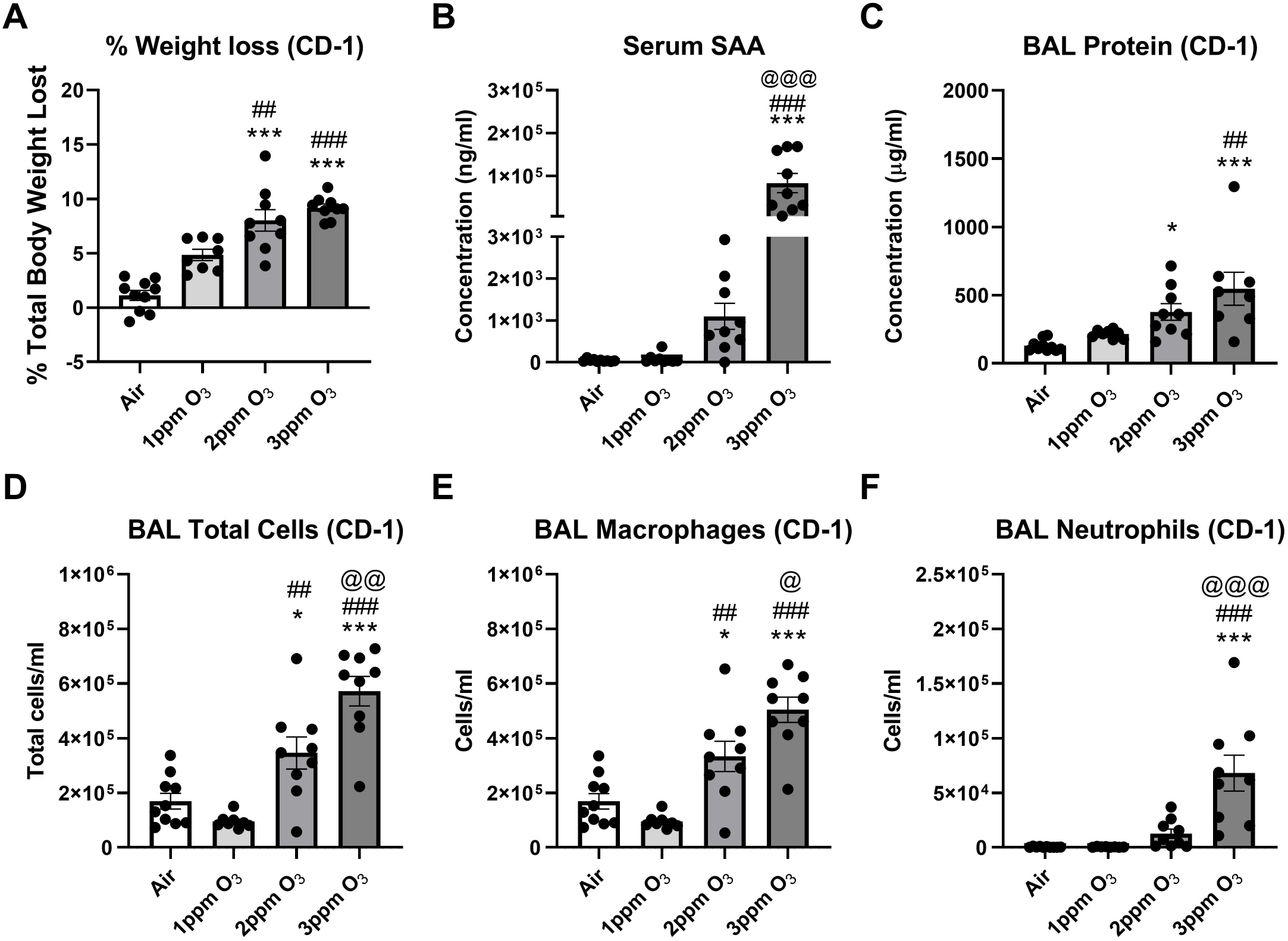
Effects of dose on O_3_-induced changes weight loss (A), serum SAA (B), BAL total protein (C), and cellular markers of acute pulmonary inflammation (D-F) in female CD-1 mice. All mice were studied 24 hours post-exposure. N= 6-10/group, *p<0.05, ***p<0.001 vs. air; ##p<0.01, ###p<0.001 vs. 2ppm; @p<0.05, @@p<0.01, @@@p<0.001 vs. 1ppm.

**Table 3.** Summary of significant dose-dependent changes induced by O_3_ in female CD-1 mice. ↑ and ↓ indicate an increase or decrease vs. air, respectively. An * indicates that the change was significantly different from the next lowest dose (i.e. 3ppm vs. 2ppm or 2ppm vs. 1ppm). N.S. = no significant difference with O_3_, N.D. = not determined.

|  |  | 1ppm O<sub>3</sub> | 2ppm O<sub>3</sub> | 3ppm O<sub>3</sub> |
| --- | --- | --- | --- | --- |
| **Pulmonary inflammation/damage markers** | Bal protein | N.S. | ↑ | ↑ |
|  | Total BAL cells | N.S. | ↑* | ↑* |
|  | BAL macrophages | N.S. | ↑* | ↑* |
|  | BAL neutrophils | N.S. | N.S. | ↑* |
| **Systemic inflammatory markers** | Serum SAA | N.S. | N.S. | ↑* |
| **Sickness Behaviors** | Weight loss | N.S. | ↑* | ↑ |
|  | Food intake | ↓ | ↓ | ↓ |
|  | Open field activity | N.S. | ↓ | ↓ |
|  | Sucrose preference | N.S. | ↓* | ↓ |

### Time-response effects of O_3_ on body weight, serum SAA, and pulmonary damage and inflammation

We next evaluated how long serum SAA increases persist, and associated changes in weight loss and pulmonary inflammation over time after a 3ppm O_3_ exposure. In this cohort, all mice were female Balb/c and were exposed to O_3_ or air at the same time to ensure uniformity of the exposures. All mice were compared to a 72-hour air control group to minimize number of animals used. Weight loss after 24 hours was slightly higher in this cohort vs. dose-response studies, with O_3_-exposed mice losing 7.6% of their initial body weight after 24 hours, which was significantly different vs. air controls. Mice gained back some of their lost weight after 48 and 72 hours, and net weight loss was significantly less at these time points vs. 24 hours, but remained significantly higher than air control (Figure 4A). SAA levels in serum were significantly elevated at 24 hours to similar concentrations observed in the dose-response cohort. SAA concentrations were significantly reduced in blood by 48 hours post-exposure, and although the mean was still arithmetically higher than the air control, there was not a statistically significant difference. Serum SAA concentrations returned to air control levels by 72 hours post-exposure (Figure 4B). Significant elevations in BAL total protein were observed up to 48 hours post-exposure (Figure 4C), whereas significant elevations in BAL total cells and neutrophils were only significantly increased vs. air control at 24 hours (Figure 4 D and F). BAL macrophages showed an approximate doubling in numbers at all time points vs. air controls, but the increases were not statistically significant vs. air (Figure 4E).

**Figure 4:**
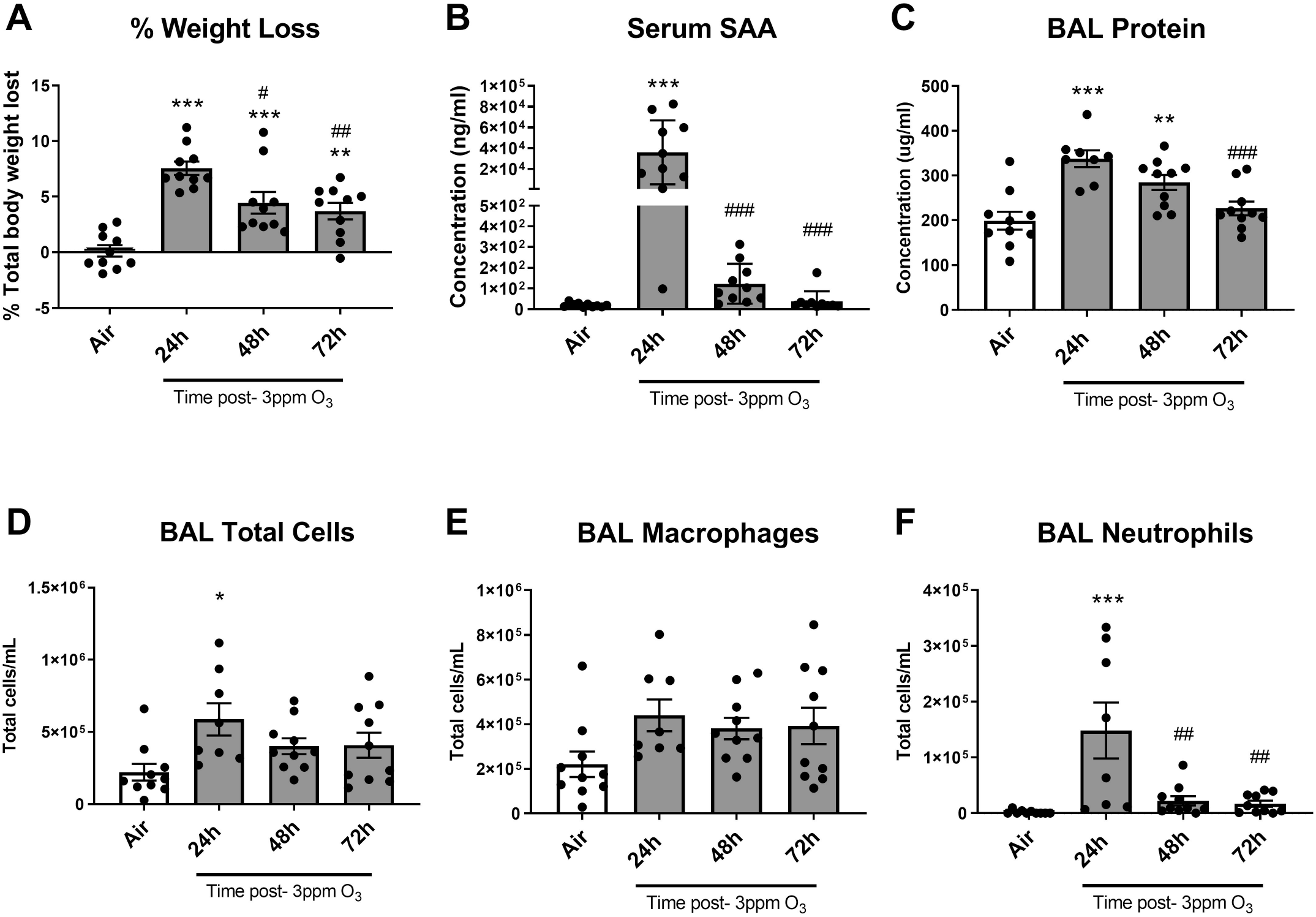
Effects of time after 3ppm O_3_ exposure on weight loss (A), serum SAA (B), BAL total protein (C), and cellular markers of acute pulmonary inflammation (D-F). The air group was studied 72 hours post-exposure. N= 8-10/group, *p<0.05, **p<0.01, ***p<0.001 vs. air; #p<0.05, ##p<0.01, ###p<0.001 vs. 24 hours.

### Effects of sex on O_3_-induced changes in body weight, serum SAA, and pulmonary damage and inflammation

Sex differences in pulmonary responses to acute ozone exposures have been reported, with young females generally showing heightened inflammatory responses in the lungs and males showing greater airway hyperresponsiveness ^51,52^. Considering that other physiological responses to O_3_ may vary by sex, we determined whether sex influenced SAA responses and associated changes in weight loss and pulmonary inflammation 24 hours following a 3ppm exposure to O_3_ in Balb/c mice. There was a significant interaction between the effects of sex and treatment on body weight loss (F(1,36)=8.915, p=0.0051), and main effects of sex and treatment on body weight loss were also significant (F(1,36)=4.464, p=0.0416 and F(1,36)=201.3, p<0.0001, respectively). The average weight of male mice prior to O_3_ exposure was 24.07 ± 1.23g, and the average weight of females was 18.95 ± 1.15g. Multiple comparisons testing showed significant differences in means of males and females exposed to air vs. O_3_, and also a significant difference in mean % body weight lost in O_3_-exposed males vs females, with females showing more weight loss than males (Figure 5A). There was a significant interaction between the effects of sex and treatment on SAA levels in serum (F(1,36)=5.784, p=0.0214), and main effects of sex and treatment on serum SAA were also significant (F(1,36)=5.785, p=0.0214 and F(1,36)=30.78, p<0.0001, respectively). Multiple comparisons testing showed significant differences in means of females exposed to air vs. O_3_, and also a significant difference in serum SAA levels in O_3_- exposed males vs. females, with females showing higher levels of serum SAA post-O_3_ exposure (Figure 5B). There was no significant interaction or main effect of sex on total BAL protein, but there was a significant main effect of treatment (F (1,33)=50.59, p<0.0001). Multiple comparisons testing showed significant differences in means of males and females exposed to air vs. O_3_, but no significant difference in means for O_3_-exposed males vs. females (Figure 5C). There was a significant interaction between the effects of sex and treatment on total BAL cells (F(1,36)=6.088, p=0.0185), and main effects of treatment were also significant (F(1,36)=52.87, p<0.0001). Multiple comparisons testing showed significant differences in means of males and females exposed to air vs. ozone, and also a significant difference in total BAL cells in O_3_-exposed males vs. females, with females showing a larger increase vs. males (Figure 5D). A significant interaction was observed between the effects of sex and treatment on BAL macrophages (F(1,36)=8.587, p=0.0058), and main effects of sex and treatment were also significant (F(1,36)=5.779, p=0.0215 and F(1,36)=41.86, p<0.0001, respectively). Multiple comparisons testing showed significant differences in means of females exposed to air vs. ozone, and also a significant difference in mean BAL macrophages in O_3_-exposed males vs. females, with females showing greater increases in BAL macrophages vs. males (Figure 5E). A significant interaction was observed between the effects of sex and treatment on BAL neutrophils (F (1,36)=7.004, p=0.0120), and main effects of sex and treatment were also significant (F(1,36)=7.372, p=0.0101 and F(1,36)=45.22, p<0.0001, respectively). Multiple comparisons testing showed significant differences in means of males and females exposed to air vs. O_3_, and also a significant difference in mean BAL neutrophils in O_3_-exposed males vs females, with females showing greater increases in BAL neutrophils vs. males (Figure 5E). These observed sex differences are summarized in Table 2.

**Figure 5:**
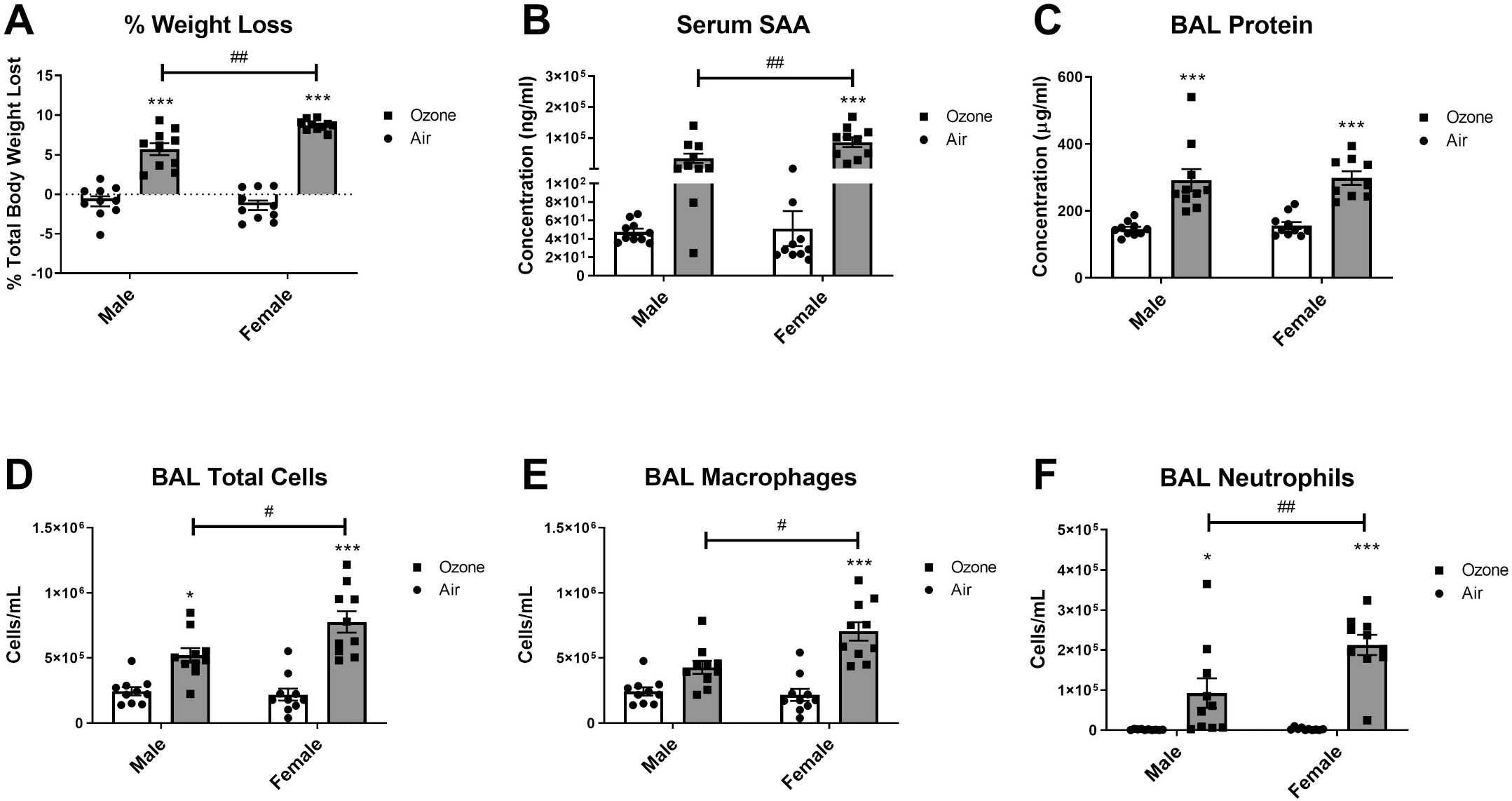
Effects of sex on weight loss (A), serum SAA (B), BAL total protein (C), and cellular markers of acute pulmonary inflammation (D-F) following 3ppm O3 exposure. All groups were studied 24 hours post-exposure. N= 9-10/group, *p<0.05, ***p<0.001 vs. air; #p<0.05, ##p<0.01 vs. groups indicated.

### Dose, time, and sex-dependent effects of O_3_ on sickness and depressive-like behaviors

Acute inflammatory insults can cause behavioral changes that include reduced food and fluid intake, weight loss, reduced locomotor activity, anhedonia, and other behavioral sequalae that have many similarities with depression ^38^. We first evaluated dose-, time-, and sex-dependent changes in food intake among mice exposed to O_3_. The doses and time points used were chosen to match the time points that were used to measure SAA and pulmonary leakage/inflammation, which was maximal at 24 hours. We found that a 1ppm O_3_ exposure caused about a 25% reduction in food intake vs. baseline at 24 hours post-exposure in female Balb/c mice (Figure 6A). A 3ppm O_3_ exposure caused a larger decrease in food intake in Balb/c female mice of about 70% (Figure 6B) to 90% (Figure 6C). Decreases in food intake in female mice persisted for 24 hours, and returned to baseline levels within 48 hours (Figure 6B). In the male-to-female comparison, there was a main effect of treatment on food intake (F(1,36)=119.5, p< 0.0001), but no significant effect of sex. Multiple comparisons testing also revealed a significant difference between air and O_3_ exposure within sexes, but food intake in male and female O_3_-exposed mice was not significantly different (Figure 6C). Prior to evaluating sucrose preference, we addressed the possibility that O_3_ may damage olfactory epithelium, causing olfactory dysfunction which could confound behavioral tests that rely on the sense of smell or taste. We did not specifically investigate whether taste perception is altered, because tests used to evaluate taste perception are likely to be confounded by changes in taste preference that are known to occur as a component of sickness and depressive-like behavioral responses to inflammatory stimuli (78). Further, mice are obligate nasal breathers and so the majority of inhaled O_3_ would be encountered in the upper airway vs. the mouth. Our data show that the ability to locate a buried hidden treat was not altered in female mice exposed to 3ppm O_3_ for 4 hours vs. air-exposed mice (Figure 6D), indicating that the mouse’s ability to detect sucrose in their drinking solution would not be influenced by severe olfaction deficits in O_3_-exposed mice. We next administered the sucrose preference test in mice, and reported both total fluid intake, and % of baseline preference for sucrose; baseline sucrose preference was about 80% in both sexes. Total fluid intake was acutely decreased by about 28% in 1ppm O_3_-exposed mice 24 hours after exposure, which was significantly different vs. control (Figure 7A). 3ppm O_3_ induced more substantial reductions in total fluid intake of about 59% (Figure 7B) to 61% (Figure 7C) 24 hours after exposure, which were also significantly different from air controls. Effects of O_3_ on fluid intake returned to baseline by 48 hours post-exposure (Figure 7B). Fluid intake in the male-to-female comparison showed a significant main effect of treatment on fluid intake (F (1, 36) = 196.7, p<0.0001), but there was no significant effect of sex or interaction between sex and treatment. We evaluated sucrose preference in the same cohort of mice by determining the change in sucrose preference from baseline, which was measured the night prior to O_3_ exposure. Sucrose preference significantly decreased by about 6.2% in 1ppm O_3_-exposed female mice 24 hours after exposure (Figure 7D). 3ppm O_3_ induced more substantial reductions in sucrose preference of about 24.7% (Figure 7E) to 39.6% (Figure 7F) in female mice 24 hours after exposure which were significantly different from air control. Effects of O_3_ on sucrose preference returned to baseline by 48 hours post-exposure (Figure 7E). Sucrose preference in the male-to-female comparison showed a main effect of treatment on fluid intake (F (1, 31) = 38.94, p<0.0001), but there was no significant effect of sex or interaction between sex and treatment. 3ppm O_3_ induced significant reductions in locomotor activity in female but not male mice (Figure 8A). Open field activity in the male-to-female comparison showed a significant main effect of treatment (F (1, 36) = 7.485, p=0.0096), and sex (F (1, 36) = 5.836, p=0.0209) but there was no significant interaction between sex and treatment. Neither O_3_ treatment nor sex had significant effects on tail suspension immobility time (Figure 8B). In repeating the evaluation of sickness and depressive-like behaviors in female CD-1 mice following three different O_3_ doses, it was found that there was a significant (p<0.001) linear trend relating O_3_ dose to measured parameters, which included food intake (F (1, 29) = 119.5), sucrose preference (F (1, 22) = 36.10), and open field activity (F (1, 33) = 28.03). Group mean differences were also assessed, and significant differences are shown in Figure 9 and summarized in Table 3.

**Figure 6:**
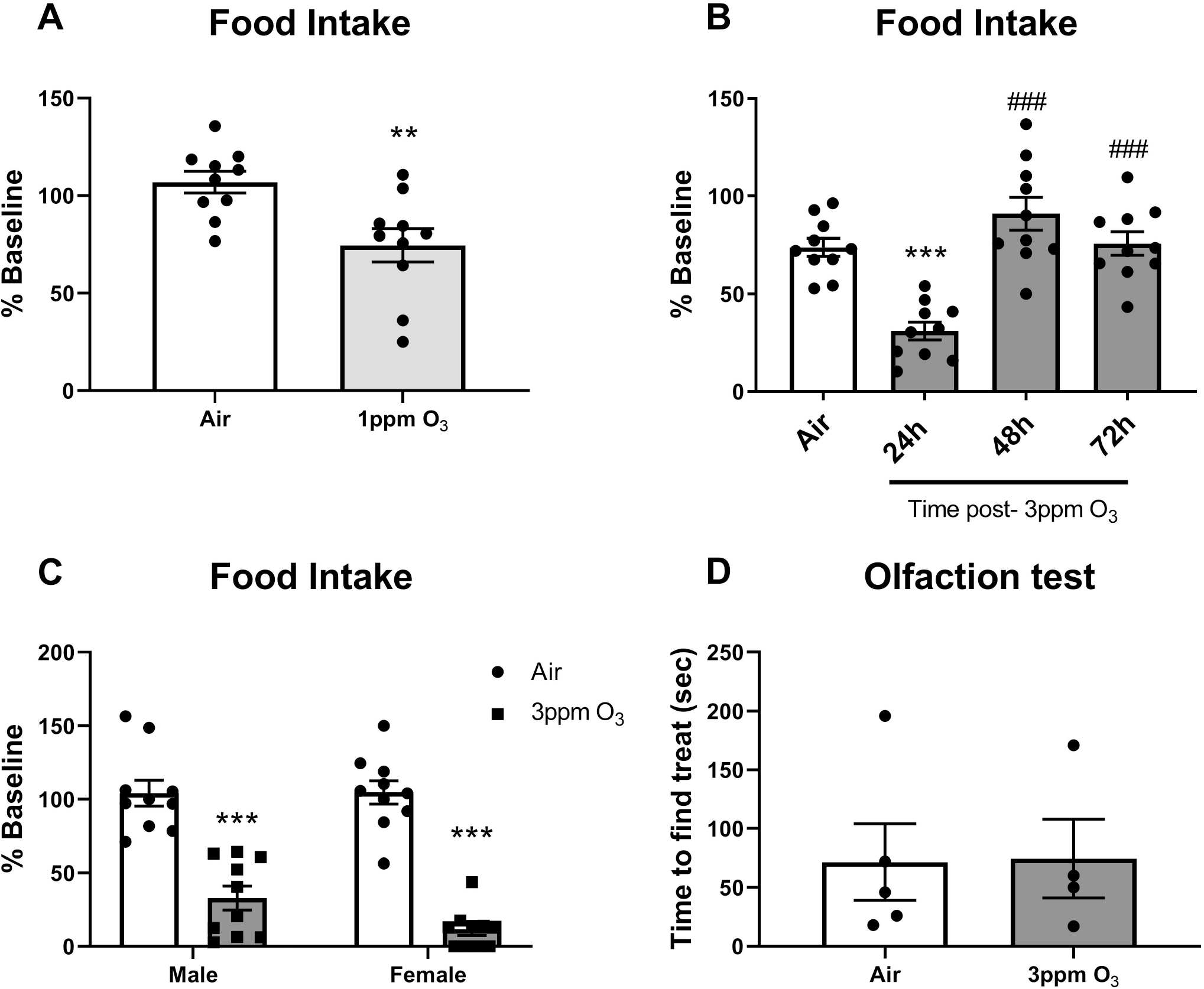
Effects of dose (A-C), time (B), and sex (C) on O_3_-induced changes in food intake, and effects of 3ppm O_3_ on olfaction (D). Except for B, all groups were studied 24 hours post-exposure. N= 10/group (A-C), **p<0.01,***p<0.001 vs. air; ###p<0.001 vs. 24h O3. N=4/group (C).

**Figure 7:**
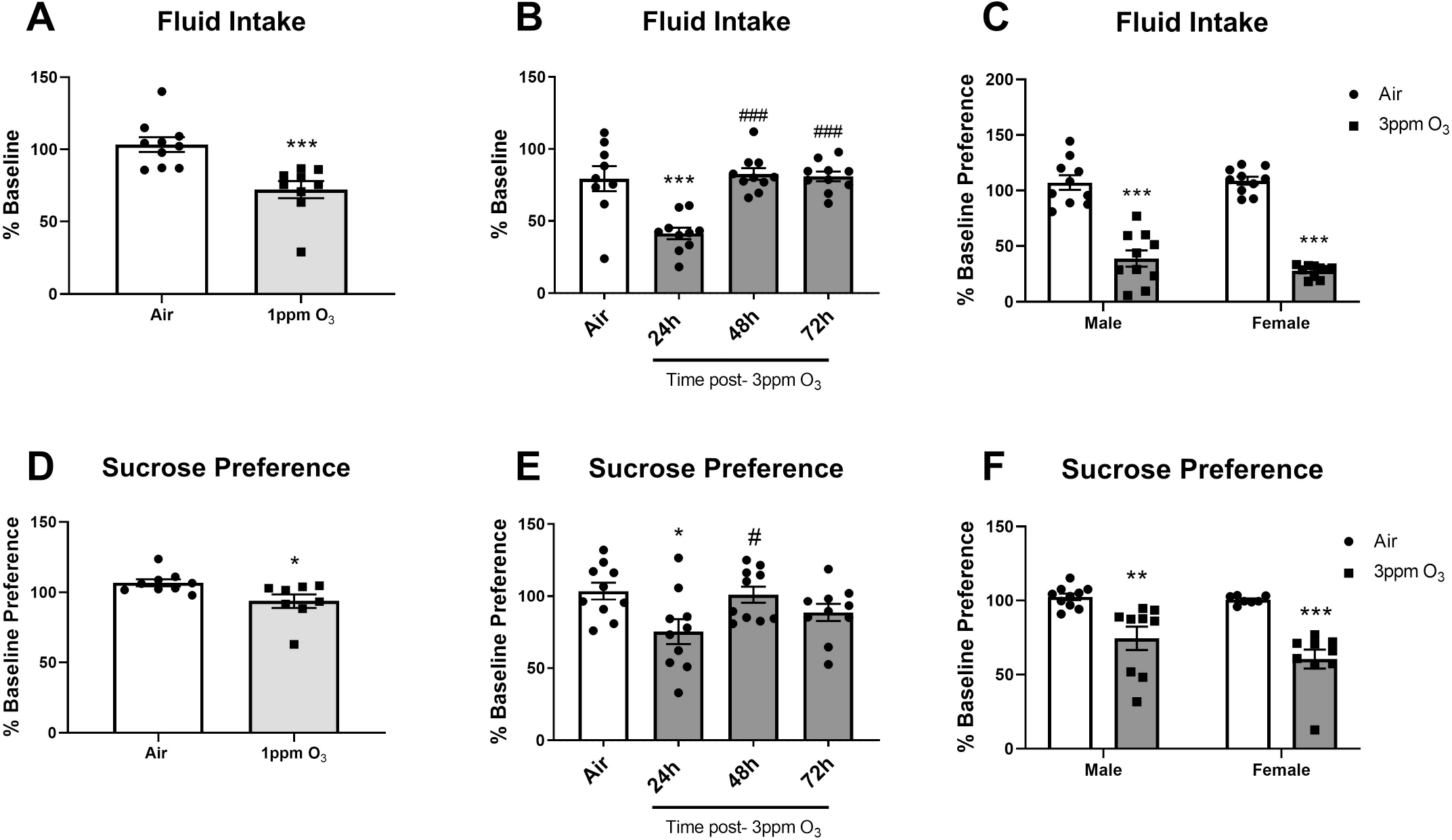
Effects of dose, time, and sex on O_3_-induced changes in total fluid intake (A-C) and sucrose preference (D-F). Except for B and E, all groups were studied 24 hours post-exposure. N= 7-10/group, *p<0.05, **p<0.01, ***p<0.001 vs. air (within sex for C and F); #p<0.05, ###p<0.001 vs. 24h O_3_.

**Figure 8:**
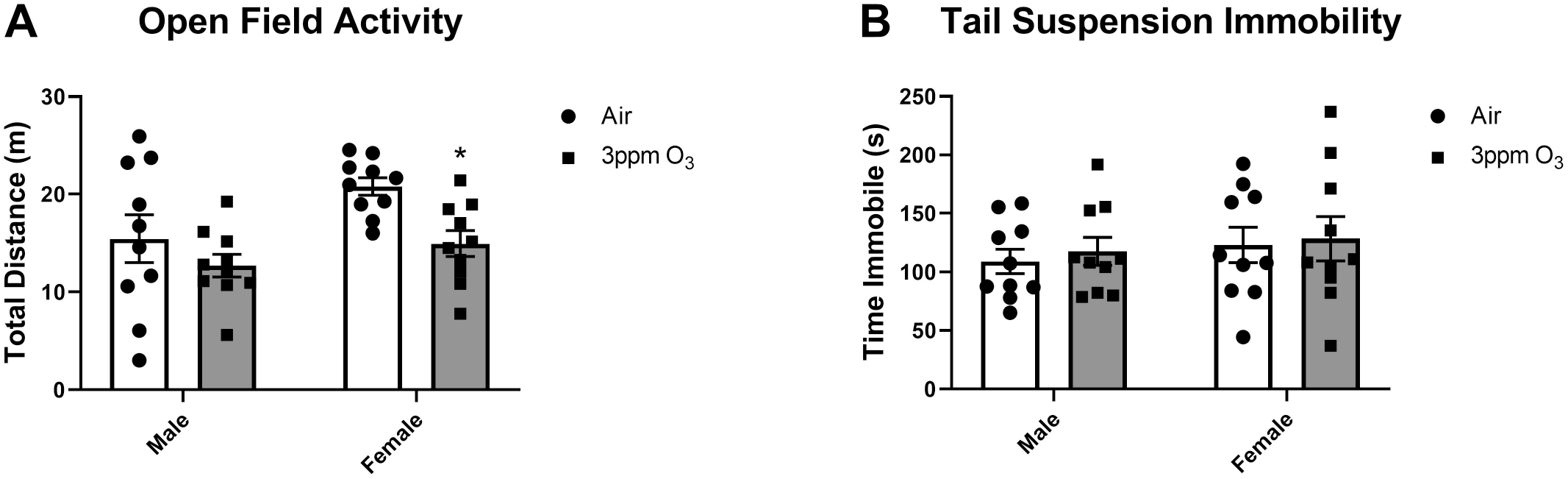
Effects of 3ppm O_3_ on open field activity (A) and immobility on the tail suspension test (B). All groups were studied 24 hours post-exposure. N= 9-10/group, *p<0.01 vs. air compared within sex.

**Figure 9:**
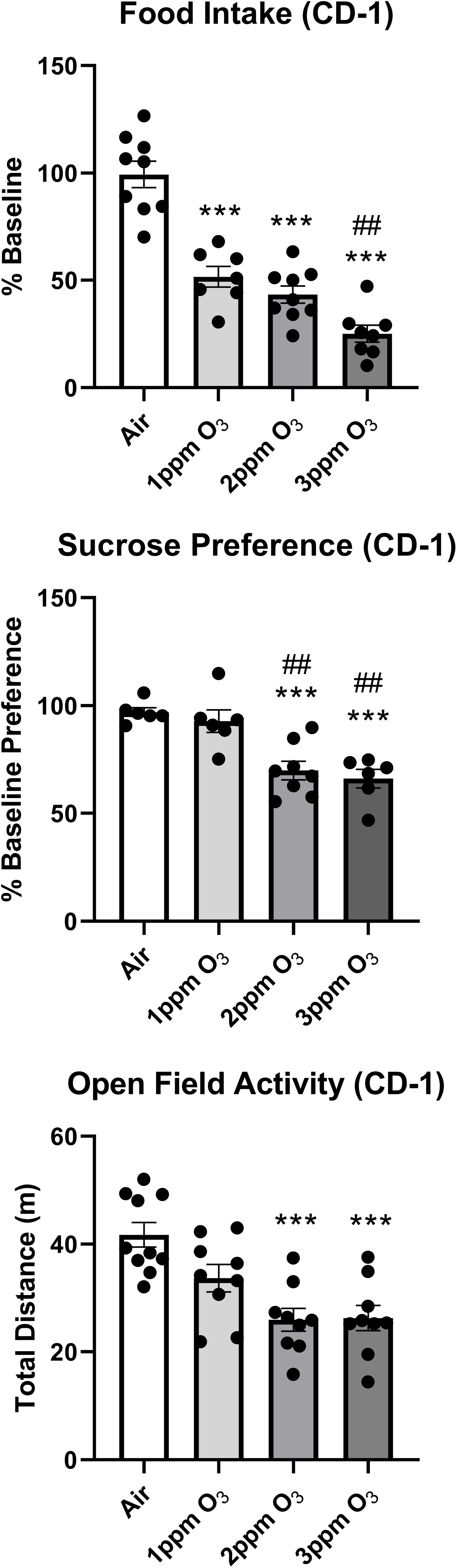
Effects of dose on O3-induced changes weight loss (A), serum SAA (B), BAL total protein (C), and cellular markers of acute pulmonary inflammation (D-F) in female CD-1 mice. All mice were studied 24 hours post-exposure. N= 6-10/group, *p<0.05, ***p<0.001 vs. air; ##p<0.01, ###p<0.001 vs. 2ppm; @p<0.05, @@p<0.01, @@@p<0.001 vs. 1ppm.

### Biochemical changes in the kynurenine pathway

IDO-dependent increases in circulating kynurenine have been found to mediate depressive-like behaviors but not sickness behaviors following an acute inflammatory stimulus ^29,30^. Kynurenine is a product of tryptophan metabolism via two rate-limiting enzymes: IDO and tryptophan 2,3-dioxygenase (TDO). TDO is predominantly and constitutively expressed in the liver by the Tdo2 gene ^53^. IDO is expressed predominantly in extrahepatic tissues including epididymis, intestines, spleen, lung, and brain by the Ido1 gene ^54^, and predominantly contributes to circulating kynurenine levels at baseline and during inflammatory conditions ^29,55^, although TDO can also contribute to circulating kynurenine pools ^56^. We therefore probed for *Ido1* mRNA expression in the spleen, brain, and lung (Figure 10A-C) and for *Tdo2* expression in the liver (Figure 10D). There was a significant main effect of sex (F (1, 26) = 7.317, p=0.0119) on *Ido1* expression in spleen and multiple comparisons testing showed that *Ido1* was significantly upregulated in spleens of female mice. There was a significant main effect of sex and treatment (F (1, 36) = 5.448, p=0.0253, F (1, 36) = 8.822, p=0.0053, respectively) on *Ido1* expression in brain, and multiple comparisons testing showed that *Ido1* was significantly upregulated in brains of female mice. In contrast, lungs showed a significant main effect of treatment (F (1, 36) = 10.83, p=0.0022), and was significantly downregulated in female mice. There were no significant effects of sex or O_3_ exposure on *Tdo2* expression. There was a significant main effect of sex and treatment on serum kynurenine levels (F (1, 32) = 20.93, p<0.0001,F (1, 32) = 18.38, p=0.0002, respectively). Multiple comparisons testing revealed that O_3_ induced significant kynurenine elevations in females, but not males, consistent with the *Ido1* expression results. Results for measured changes 24 hours after exposure are summarized in Table 2.

**Figure 10:**
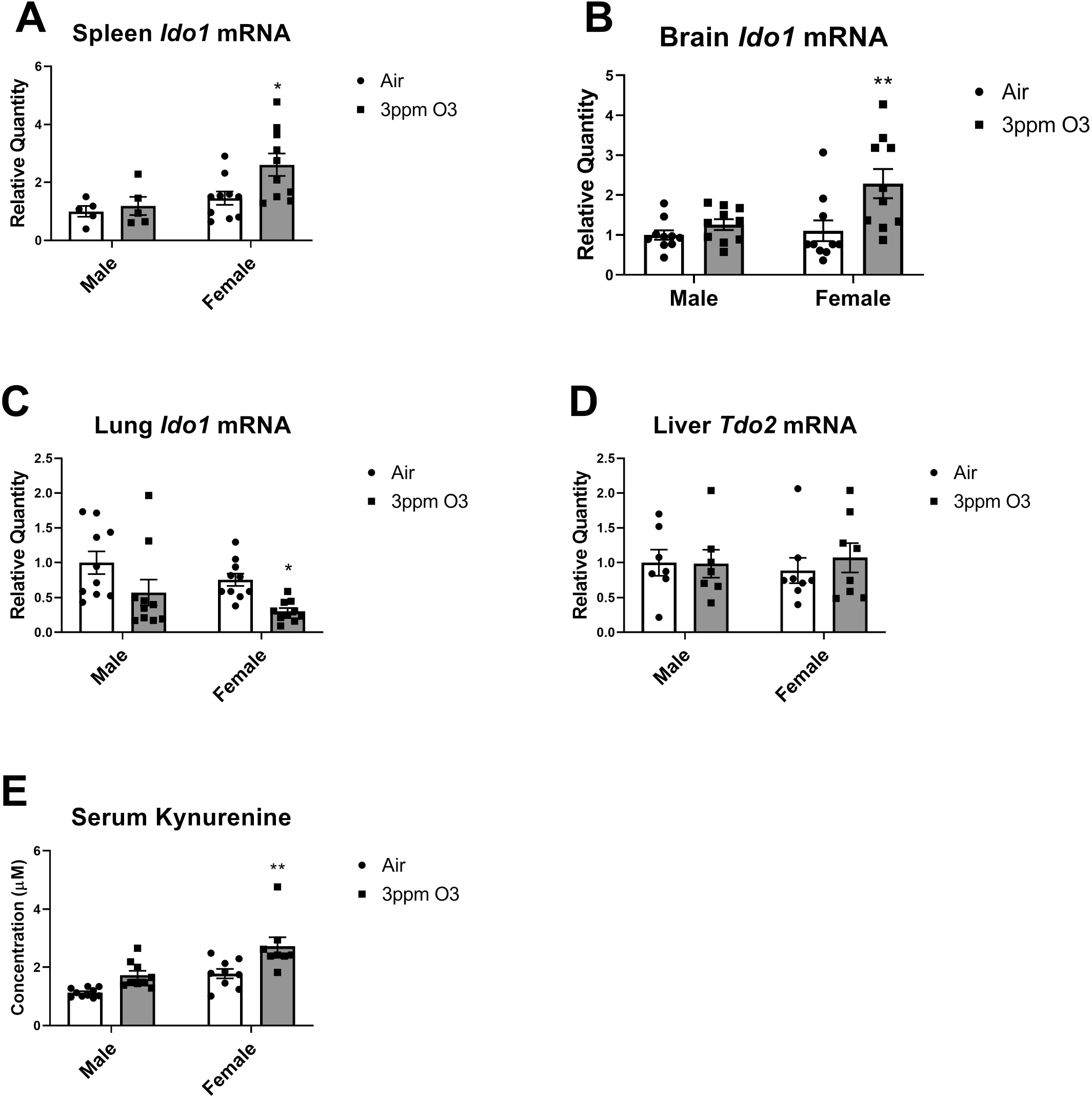
Effects of 3ppm O_3_ on *Ido1* mRNA expression in spleen, brain, and lung (A-C), *Tdo2* mRNA expression in liver (D), and kynurenine concentrations in serum (E). All groups were studied 24 hours post-exposure. N= 8-10/group, *p<0.05, **p<0.01 vs. air compared within sex.

## Discussion

Although O_3_ primarily targets the lungs, O_3_ exposure can also have effects on distal organs that include the CNS ^8,57,58^. How O_3_ alters CNS functions are incompletely understood, but recent works support a lung-brain axis mechanism whereby circulating factors are upregulated in response to pulmonary inflammation ^12,25^. These circulating factors can then exert their effects in the brain directly if they can cross the BBB, or may have indirect effects by modifying functions of brain barriers or activating regions of the brain that lack a BBB such as circumventricular organs. Two circulating factors that are induced by O_3_ and can cross the intact BBB are SAA and kynurenine. Both molecules have neuromodulatory functions, and can cause depressive-like behaviors. To better understand the relations of pulmonary inflammation, neurobehavioral changes, and circulating SAA/kynurenine, we conducted dose-response and time-response studies following a 4-hour single exposure to O_3_, and also determined whether these responses varied by sex.

Our prior work has shown that O_3_ increases production of SAA in the liver, leading to increased SAA in blood which then crosses the intact BBB ^25^. We also showed previously that SAA induction in blood is significantly correlated with cellular markers of pulmonary inflammation ^25^, indicating that SAA concentrations in blood may be related to the degree of pulmonary inflammation caused by O_3_. One limitation of this study was that it was only conducted at a 3ppm dose, which elicits a high level of pulmonary inflammation in mice relative to that observed in O_3_ -exposed humans ^6,7,43,44^. Therefore, we conducted dose-response studies at 1ppm and 3ppm in female Balb/c mice and 1, 2, and 3ppm in female CD-1 mice to estimate the effective dose needed to elicit changes in systemic biomarkers and behaviors. We showed that significant SAA elevations in blood occurred 24 hours after a 3ppm but not a 2ppm or 1ppm O_3_ exposure. Increased BAL total cell counts and neutrophil counts were also significantly elevated only at the 3ppm O_3_ dose, which supports a relation of hepatic SAA induction and release into blood with pulmonary neutrophilia. Total BAL protein, which is a marker of alveolar-capillary barrier function ^50^, was equally elevated by 1ppm and 3ppm O_3_ in Balb/c mice, and in a dose-dependent manner in CD-1 mice. The finding that BAL protein elevations occurred at a lower O_3_ dose that did not induce significant increases in SAA suggests that SAA does not contribute to alveolar-capillary barrier dysfunction in this model. However, it is possible that SAA from blood could enter the pulmonary compartment via leakage mechanisms and exert local effects. Food intake, fluid intake, and sucrose preference were also significantly reduced by both 1ppm and 3ppm O_3_ in Balb/c mice, but 3ppm O_3_-induced decrements were greater in magnitude. There was a dose-dependent relation of these parameters in CD-1 mice. These findings indicate that there is a dose-effect of O_3_ on sickness and some depressive-like behaviors, and suggest that SAA does not contribute to the behavioral effects at the 1ppm dose. One limitation of our findings is that the 3ppm O_3_ dose elicits more severe pulmonary inflammation in mice than what has been reported in humans exposed to relatively high concentrations of O_3_ vs. peak O_3_ ambient concentrations in most of the world ^43,44^, and so the relevance to human ambient exposures is currently unclear. However, there may be increased vulnerability to the harmful effects of O_3_ with chronic dosing, preexisting medical conditions, or use of O_3_ generators in personal or industrial settings that exceed OSHA standards.

Our findings on the relation of SAA induction in blood and lung neutrophilia are consistent with prior works from other groups who have showed that SAA may be both a systemic marker and mediator of pulmonary injury. For example, it has been shown that SAA levels in lungs of chronic obstructive pulmonary disease (COPD) patients positively correlate with elastase-positive neutrophils, and that increases in circulating SAA can predict severity of acute exacerbations of COPD ^34^. Further, chronic intranasal treatments of SAA induced neutrophilic airway inflammation ^59^. Mouse SAA3 may also be a mediator of IL-6-dependent pulmonary inflammation through its ability to induce IL-17A ^60^.

Time-response studies showed that O_3_’s effects on SAA levels in blood, markers of lung damage and inflammation, and measures of sickness and depressive-like behaviors were most substantial at 24 hours post-exposure, and most of the measured parameters returned to control levels by 48 hours post-exposure. Parameters affected by O_3_ which remained changed at 48-72 hours included body weights, and total BAL protein. Elevations in BAL protein persisted up to 48 hours, but then returned back to normal by 72 hours. The persistence of increased BAL protein up to 48 hours post-exposure is consistent with time-course findings reported previously ^61^, and indicates that disruption of the alveolar-capillary barrier persists even after inflammatory markers measured here have returned to baseline. Mice lost a significant amount of weight 24 hours post O_3_-exposure at 3ppm, which was partially regained by the 48- and 72-hour time points. Food intake and fluid intake also returned to normal by 48 hours post-exposure, but there was no compensatory increase above baseline, which might explain why weights did not return to baseline levels by 72 hours.

Inflammatory stimuli, a prototype being LPS, can induce behavioral responses of sickness and depressive-like behaviors that follow distinct temporal patterns of appearance or persistence. For example, a single low-dose LPS injection results in initial cytokine-induced sickness behaviors that resolve quickly, whereas depressive-like behaviors persist for a longer time period ^28,62^. Activation of c-Fos in neuroanatomical areas known to contribute to cytokine-induced sickness behaviors were also induced at 6 hours and resolved by 24, which is consistent with induction and resolution of pro-inflammatory cytokines by LPS that can activate vagal afferents that project to these regions ^63,64^. LPS increased immobility in the forced swim and tail suspension tests and also reduced sucrose preference: these behaviors persisted for at least 24 hours ^28^, aligning temporally with the activation of delta FosB in brain regions that are implicated in depressive-like behaviors, including the extended amygdala, hypothalamus, and hippocampus ^28^. Food intake post-LPS was shown to be reduced up to 28 hours and thus persisted through both phases of behavioral change ^62^. Data shown by us and others indicate that acute O_3_ exposure, which initially induces injury through oxidative damage, elicits an inflammatory and behavioral response that is distinct from a pathogen-associated insult like LPS. First, circulating pro-inflammatory cytokines are generally not increased, or only slightly increased in blood following an acute O_3_ exposure ^7,12,25,65,66^. However, circulating SAA levels increase by orders of magnitude following O_3_ exposure, and peak around 24 hours post-exposure. Since we posited that SAA is a mediator of depressive-like behaviors in this model, we selected behavioral time points for study that would adequately capture maximal induction and recovery of SAA, ranging from 24 to 72 hours. After O_3_ exposure, significant reductions in sucrose preference were observed concurrently with reductions in total fluid intake and food intake and these behaviors resolved by the second night following O_3_ exposure. Thus, resolution of behavioral effects of O_3_ occurred together with resolution of SAA and pulmonary inflammation, although at the lower 1ppm dose of O_3_, there was no induction of SAA or lung neutrophils despite more modest effects on food/fluid intake and sucrose preference. Therefore, factors other than SAA are likely contributing to the behavioral responses to O_3_.Additional behavioral evaluations of locomotion in the open field and immobility on the tail suspension test revealed a modest but significant decrease in open field activity in females but not males, and no difference on tail suspension immobility. These findings further indicated differences in O_3_ exposure vs. LPS, in that sickness behaviors appear to persist in the presence of some, but not all, depressive-like behaviors that LPS induces.

Sex differences were observed for many measured parameters following O_3_ exposure, including the magnitude of weight loss, pulmonary inflammation, and circulating levels of SAA. Our findings are consistent with other studies showing that female mice have exacerbated pulmonary inflammation in response to O_3_ ^51,52^. One limitation of this study is that we were not able to collect data on dosimetry, and so we cannot rule out sex differences in O_3_ deposition due to size disparity. However, an evaluation of O_3_ deposition in adults vs. pups showed that there was less overall O_3_ deposition in the pups, suggesting that smaller size reduces O_3_ deposition ^67^. Therefore, we would expect a size disparity to favor slightly more O_3_ deposition in male mice. Increases in circulating kynurenine and in *Ido1* mRNA levels in brain and spleen were specifically apparent in female mice, although male-female comparisons post-O_3_ were not significantly different and there was not an interaction between sex and treatment. In lungs, *Ido1* expression was arithmetically decreased in both sexes, although the difference was only statistically significant in females. The reduction of *Ido1* in lungs is consistent with increased lung neutrophilia that occurs with O_3_, since IDO negatively regulates neutrophil trafficking ^68^. Levels of liver *Tdo2* were unchanged, suggesting that *Ido1* could be a predominant mediator of the systemic increases in kynurenine. However, future work is needed to verify that increases in circulating kynurenine are IDO-dependent, and to identify the prevailing tissue that contributes to increases in circulating kynurenine. These findings also implicate the kynurenine pathway as a second systemic mediator in the lung-brain axis, since kynurenine can cross the intact blood-brain barrier via the large neutral amino acid transporter ^32^, and the majority of brain kynurenine is derived from blood under physiological conditions. In inflammatory states, nearly all kynurenine in the brain comes from blood ^69^. In the brain, kynurenine can be metabolized to neuroprotective or neurotoxic mediators in the brain a cell-type dependent manner. In the healthy brain, kynurenine is predominantly metabolized by astrocytes which produce kynurenic acid, which is neuroprotective at physiological levels ^70^. Microglia express enzymes that metabolize kynurenine into neurotoxic products such as the radical-generating 3-hydroxykynurenine and quinolinic acid, which can cause excitotoxic cell death ^30^. Under inflammatory conditions, kynurenine, 3-hydroxykynurenine, and quinolinic acid are increased in the brain ^62^ and these increases have been proposed to contribute to neurological diseases like Alzheimer’s and depression ^31^. Our findings corroborate recent studies in rats which showed that acute O_3_ exposure increases circulating levels of kynurenine, and in this model, kynurenine upregulation was not significantly affected by the glucocorticoid inhibitor metyrapone ^26^. In another study that investigated effects of maternal O_3_ exposures and/or high-fat diet on the metabolomes of male and female offspring in rats, it was found that maternal exposure to O_3_ significantly increased the circulating kynurenine concentrations in female but not male offspring. Male offspring kynurenine levels were arithmatically higher than their female counterparts born from air or O_3_-exposed dams, but unaltered by maternal O_3_ exposure ^71^. Together, these findings implicate the kynurenine in different O_3_ exposure settings, and future work is needed to evaluate how systemic elevations in kynurenine might affect CNS functions in context of O_3_ exposures. Our prior work has shown that 3ppm O_3_ does not cause increases in brain or blood cytokines ^25^, suggesting that the inflammatory links of O_3_ to CNS dysfunction do not involve classical pro-inflammatory cytokine-mediated responses such as those to pathogen-associated stimuli.

In conclusion, our work shows that O_3_ exposure resulting in significant lung inflammation is translated to alterations in systemic inflammation, including effects on behavior. There are sex-dependent differences in the systemic effects of O_3_ exposure that correspond with those found for pulmonary inflammation, suggesting that female mice mount a more potent inflammatory response to O_3_. This study also supports that the kynurenine pathway is a possible mediator of ozone’s effects on the CNS.

